# FTO (fat-mass and obesity-associated protein) deficiency aggravates age-dependent depression-like behaviors and cognitive impairment

**DOI:** 10.1101/2024.12.04.626872

**Authors:** Mengdie Li, Yating Yang, Tangcong Chen, Yueyang Luo, Yingqian Zhang, Huanzhong Liu, Michael Maes

## Abstract

**Background:** The demethylase fat mass and obesity-related protein (FTO) is strongly associated with depression. Aging is a risk factor for synaptic plasticity damage in the brain and leads to neurocognitive dysfunctions. However, whether FTO is associated with susceptibility to depression in different age groups remains unknown.

**Methods:** We subjected 3-and 12-month-old C57BL/6J male mice to 6 weeks of chronic unpredictable mild stress (CUMS) and 3 weeks of hippocampal injection of FTO knockdown adeno-associated virus 9 shRNA (FTO-KD AAV9). Finally, 36 male mice in each 3-month-old and 12-month-old groups were divided into three groups (n=12): Sham, CUMS, and FTO-KD. After 6 weeks, we assessed behavioral deficits (depressive and anxiety-like behaviors and cognitive impairment) by behavioral tests and hippocampal neuronal damage (dendritic spine density, neuronal atrophy, and expression of proteins associated with synaptic plasticity) by molecular biochemical experiments.

**Results:** The results showed that 12-month-old C57BL/6J mice were more likely to develop depression-like behavior and spatial learning and memory impairment induced by CUMS than 3-month-old mice. Chronic stress-induced depression-like behavior and cognitive impairment worsened after the FTO-KD intervention. In the hippocampus of 3-and 12-month-old mice, CUMS induced the downregulation of FTO, nerve growth factor (NGF), reelin, and synaptic plasticity-related proteins. It also caused abnormal BDNF-TrkB signaling, reduced density of dendritic spines, and an increased number of neuronal pyknotic nuclei, leading to neuronal disarray, which was more significant in 12-month-old animals. FTO deficiency accelerated neuronal damage in the hippocampus of 12-month-old CUMS mice.

**Conclusions:** This study provides rodent evidence that FTO deficiency may increase the susceptibility to depression in older adults by impairing hippocampal neuronal function and neuronal synaptic plasticity in an age-dependent manner. This suggests that the development of FTO activators may be an effective treatment for depression in older adults.

## Introduction

Major depressive disorder (MDD) is a severe psychiatric condition that adversely affects individuals’ emotional and physical well-being. The WHO reports that over 280 million individuals globally experience depression, with around two-thirds exhibiting cognitive deficits [1–3]. Furthermore, by 2030, MDD is projected to be the leading cause of disease burden globally [4].

A study revealed that 94% of depressed patients display neurocognitive impairments after the alleviation of their depressive symptoms [5]. Cognitive deficits in depression are significantly correlated with age [6]. Neurocognitive deficits endure longer in older persons who have experienced depression and subsequently recovered, compared to younger individuals who recover from depression [5]. This suggests that the interaction between aging and depression may result in more pronounced neurocognitive deficits.

The hippocampus is a crucial brain region that requires examination of the factors contributing to cognitive impairment linked to depression. Neuroimaging and postmortem brain studies [8, 9] have demonstrated that the hippocampus tissue volume in individuals with MDD is reduced compared to that of healthy individuals and that the hippocampal morphology is changed [7, 8]. Moreover, the hippocampus is a significant area susceptible to age-related degeneration [9]. Neurotrophins have a crucial role in regulating neuroplasticity, neuroprotection, and brain development. For example, nerve growth factor (NGF), the first neurotrophic factor identified in 1951, is crucial for facilitating the growth, differentiation, and survival of neurons. Brain-derived neurotrophic factor (BDNF) is a multifunctional member of the neurotrophic factor family, extensively and abundantly expressed in many brain areas, particularly in the subventricular zone and dentate gyrus [10, 11]. Preclinical studies [12, 13] have demonstrated that NGF and BDNF are significantly concentrated in the hippocampus and prefrontal cortex, where they play crucial roles in axon growth, neuronal survival, differentiation, neuroprotection, neurogenesis, and synaptic plasticity, including postsynaptic density protein 95 (PSD-95), synaptophysin (SYN), and synaptic morphology. NFG and BDNF expression levels have been documented to be diminished in the hippocampus of individuals with mental disorders including MDD [14–16]. Rodents subjected to chronic stress exhibited reduced levels of NGF and BDNF expression in the hippocampus [17]. This indicates that the aberrant expression of NGF and BDNF is a significant pathophysiological characteristic of MDD. Reelin, an extracellular matrix protein with a molecular weight of approximately 400 kDa, is mostly secreted by γ-aminobutyric acid (GABA) neurons in the adult cerebral cortex and hippocampus, and is intricately associated with brain development [18, 19]. Hippocampal reelin facilitates dendritic spine growth and maturation, augments synaptic plasticity and hippocampus neurogenesis, and is involved in learning and memory [20, 21].

Epigenetic changes significantly contribute to the etiology of MDD. N6-methyladenosine (m6A) is a widespread mRNA methylation alteration, influencing over 7000 mRNAs in over 25% of mammalian species [22, 23]. This alteration is significantly enriched in the brain, with levels that are double those found in other tissues. Epigenetic alterations are regarded as significant indicators of cerebral and cellular aging. m6A methylation alteration plays a dynamic role in neuronal development and the aging process of the brain. Research involving humans and animals has demonstrated that m6A methylation sites markedly increase with age in the central nervous system (CNS) [24]. Fat mass and obesity-associated protein (FTO) is a crucial regulator of the dynamic equilibrium of m6A methylation modification and is significantly concentrated in the brain [25]. FTO-dependent m6A demethylation occurs in the axons and dendritic axes of neurons, indicating that the dynamic removal of m6A methylation may be crucial for brain function [26–28]. Additionally, FTO mRNA substrates include Bdnf, Myc, Jun, Olig2, and Grin1. FTO deficiency correlates with neural developmental abnormalities, growth retardation, cognitive impairment, and anxiety-and depression-related behaviors in mammals [29–31].

Nonetheless, the impact of FTO reductions and aging on stress-induced depression-like behaviors and the underlying processes remains ambiguous. Hence, this study aims to examine the impact of age-related variations in FTO on stress-induced depressive behaviors and cognitive impairments. To achieve this objective, we subjected 3-and 12-month-old C57BL/6J mice to chronic unpredictable mild stress (CUMS). FTO knockdown adeno-associated virus 9 shRNA (FTO-KD AAV9) was employed to reduce hippocampal FTO levels in order to examine its impact on depression-like behaviors and cognitive deficits in CUMS animals. We hypothesized that FTO deficiency exacerbates depressive-like behaviors and cognitive deficits in 12-month-old mice by downregulating the expression of NGF and reelin proteins in the hippocampus, augmenting neuronal damage, diminishing the expression of synaptic plasticity proteins, and inhibiting the BDNF-TrkB signaling axis.

## Materials and Methods

### Animals

Male C57BL/6J mice, aged 3 and 12 months, were supplied by Henan Skobes Biotechnology Co., LTD. Mice were housed in an environment with regulated temperature (22 ± 1 LJ), humidity (50 ± 10%), and a 12-hour light/dark cycle. Food and water were accessible at all times, 24 hours a day. All experimental protocols received approval from the Animal Ethics Committee of Chaohu Hospital, affiliated with Anhui Medical University (KYXM-202112-010).

### Groups

Following a two-week adaption phase, all mice were randomly allocated into three age-based groups: (1) 3-month-old: Sham group; CUMS group; FTO knockdown (FTO-KD) group; (2) 12-month-old: Sham group; CUMS group; FTO-KD group. During weeks 1 to 6, all groups of mice, excluding the Sham group, were subjected to one or two distinct stressors daily. In the third week, all mice were treatment via stereotactic brain surgery. All mice in the Sham and CUMS groups were administered enhanced green fluorescent protein (EGFP) AAV9 shRNA, while all animals in the FTO-KD groups were administered FTO-KD AAV9 shRNA (Shanghai Genechem Co., Ltd, China). Stressors, such as food or water deprivation, were only sparingly applied within one week of surgery. At the conclusion of week 6, behavioral assessments were conducted for a duration of 2 weeks, after which the mice were euthanized immediately following the experiment. The body weights of all mice were documented at 0, 2, 4, and 6 weeks throughout the investigation. **Figure 1** illustrates the experimental methodology and CUMS stressors.

**Figure 1.**
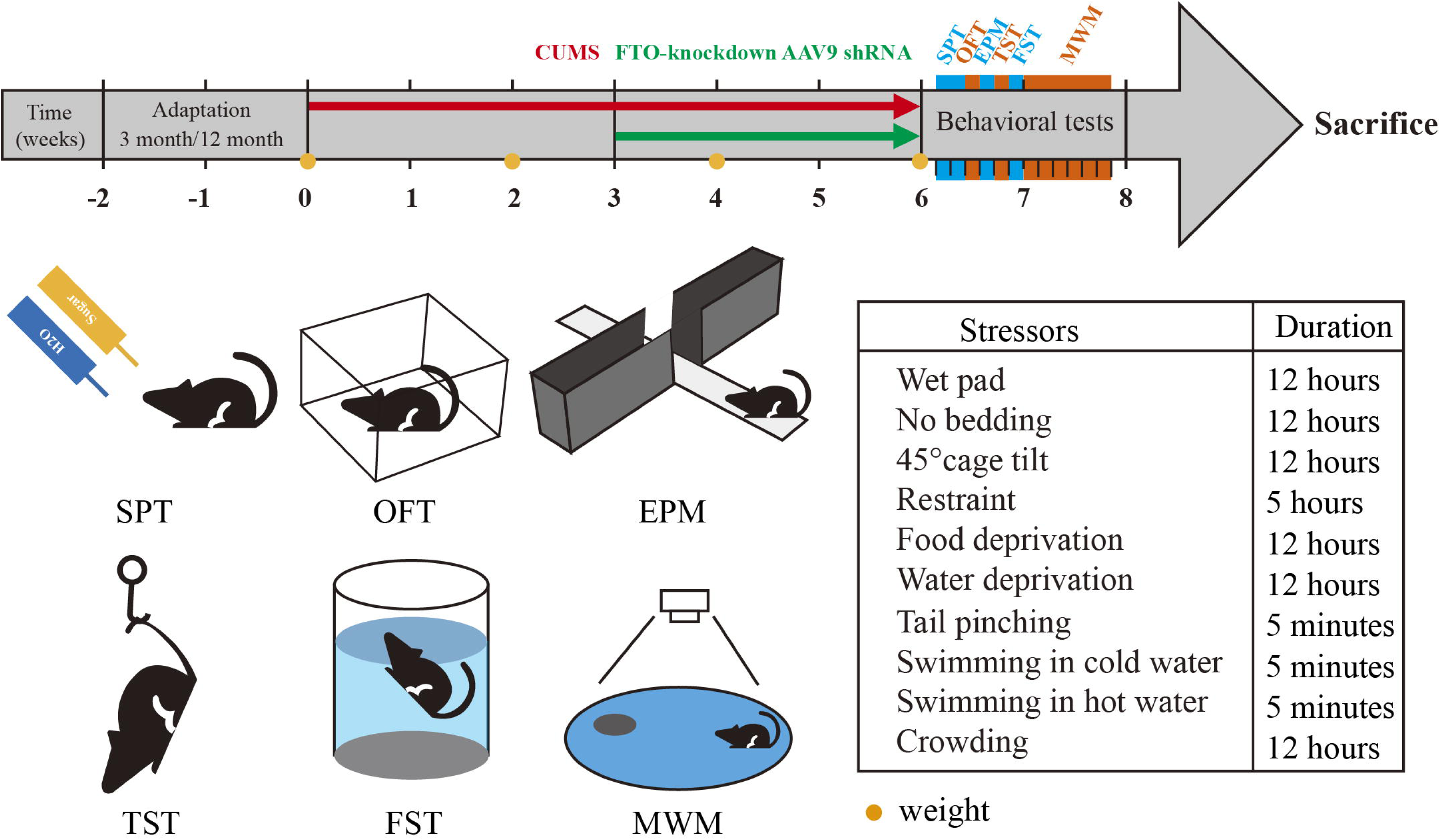
Experimental design and protocol. The CUMS protocol includes 10 stressors. CUMS: Chronic Unpredictable Mild Stress; AAV9: adeno-associated virus 9; SPT: Sucrose preference test; OFT: Open field test; EPM: The elevated plus maze; TST: Tail suspension test; FST: Forced swimming test; MWM: Morris Water Maze.

### Stereotaxic Surgery

The stereotactic surgical apparatus was supplied by Reward Life Technology Co., LTD (Shenzhen, China). The injection site was located in the bilateral hippocampus: anteroposterior = -2.1 mm, mediolateral = ± 2.1 mm, dorsoventral from the bregma = -2.0 mm. Each side received an injection for 4 minutes at a rate of 250 ng/min, totaling 4 µg, with the needle retained for an additional minute, culminating in a total duration of 5 minutes. The vital signs of the mice were meticulously monitored post-surgery.

## Behavior tests

### Sucrose preference test (SPT)

SPT was employed to identify anhedonia in animals, and we made minor enhancements to a prior experimental technique [32]. Prior to behavioral testing, all mice were individually housed in cages of 15 × 15 × 15 cm³, with one cage equipped with two bottles. On the initial day, all mice were provided with two water bottles and maintained for 24 hours. The following day, one of the bottles was substituted with a 1% sucrose solution, and the mice had unrestricted access for 12 hours. The bottles containing water and sucrose were subsequently switched and administered for an additional 12 hours. On the third day, the mice were denied access to water and food for a duration of 24 hours. On the fourth day, each mouse received two pre-measured solutions: 120 mL of water and 120 mL of 1% (w/v) sucrose solution. After four hours, the bottles were weighed and repositioned, and after a further four hours, they were weighed once again. Sucrose preference ratio (%) = [sucrose consumption (mL) / (sucrose consumption (mL) + water consumption (mL))] × 100%.

### Open field test (OFT)

The OFT is employed to observe the spontaneous locomotion of animals in an open field setting [21]. Prior to the behavioral assessment, all mice were adapted to the testing environment for a duration of 24 hours. Initially, mice were permitted to explore freely in the open field for 10 minutes, with their activity quantified via the RWD automatic tracking system (RWD, Shenzhen, China), encompassing both the central (25 × 25 cm²) and peripheral regions (50 × 50 cm²).

### Elevated-plus maze (EPM) test

This study employed the EPM test to evaluate anxiety-like behavior in animals [33]. The apparatus comprises two enclosed and two open arms, positioned 50 cm above the ground. Initially, the mice were positioned at the center of the maze, oriented towards the open arm. A video tracking system (RWD, Shenzhen, China) was employed to assess and document murine behavior. The distance and retention time of the mice in the open arm were evaluated throughout a 5-minute period.

### Tail suspension test (TST)

The TST was employed to assess depression-like behavior in animals, as described by prior research [34]. The mouse’s tail was secured to a hook with tape, orienting it upside down, while its head was positioned 15 cm above the ground. The experimental area consisted of a black wooden frame. A high-definition camera filmed the mice’s behavior for 6 minutes, comprising a 2-minute latency phase followed by a 4-minute test period, during which the immobility time of the mice was documented (RWD, Shenzhen, China).

### Forced swimming test (FST)

The FST was conducted in accordance with experimental methods described previously [35]. A transparent cylindrical bucket was filled with water at a temperature of 23 ± 1 LJ to a depth of 15 cm. A camera tracking system (RWD, Shenzhen, China) was employed to document the swimming behavior of the mice in water for 10 minutes, and the duration of immobility (s) of the mice was recorded. Upon completion of the experiment, the mice were dried using a heater and subsequently returned to their habitat.

### Morris Water Maze (MWM)

The MWM was employed to evaluate spatial learning and memory [36]. The MWM was conducted in a 60 cm deep white circular pool at a temperature of 22 LJ ± 1. Mice underwent four training trials daily for five consecutive days to locate platforms utilizing information from surrounding visual cues. During the initial five days, the mice were allotted 60 seconds to locate the platform. If the mice failed to locate the platform, they were aided in finding it and remaining on it for 30 seconds. On day six, the platform was removed, and the mice were allotted 60 seconds to locate the hidden platform. The camera (RWD, Shenzhen, China) documents the frequency of entries into the platform, duration spent in the target quadrant, swimming velocity, and the count of entries into the target quadrant, while the Smart program was employed to assess these metrics.

## Assays

### Immunohistochemistry (IHC)

The expressions of hippocampus FTO, NGF, and reelin were identified following a previously established experimental methodology [37]. The antibodies employed were rabbit anti-FTO (Proteintech, 1:500), rabbit anti-NGF (Affinity, 1:100), and rabbit anti-Reelin (Abcam, 1:300). The secondary antibodies utilized were HRP goat anti-rabbit (Servicebio, 1:200).

### Hematoxylin-eosin (H&E) staining

H&E staining was conducted with a previously published method [38]. Following 4% paraformaldehyde perfusion, the entire brain was dissected. The hippocampus was fixed in paraffin and sectioned to a thickness of 5 µm. Dewaxing was performed with xylene, and rehydration was conducted with 100% ethanol. Hematoxylin was applied for 5 minutes, followed by eosin counterstaining for 3 minutes. Subsequently, it was processed with ethanol, xylene, and resin. H&E staining was employed for morphological examination utilizing a high-resolution optical microscope (Olympus, Japan).

### Golgi-cox staining

Golgi-Cox staining was conducted in accordance with prior research [25]. Following the deep anesthesia of the mice, the entire brain was swiftly excised. The entire brain was sectioned into tissue blocks around 10 mm in thickness. Brain tissue specimens were subjected to staining using Golgi staining solution (Servicebio, Wuhan, China). The specimens were immersed in the staining solution, which was replaced after 48 hours, and subsequently treated in darkness for 14 days, with the staining solution being changed every 3 days. Sections of 60 µm in thickness were excised utilizing a vibrating microtome (Leica, Germany).

### Western blotting

Proteins were identified using Western blotting as previously outlined. Rabbit anti-FTO (1:5000, Thermo, USA), rabbit anti-BDNF (1:1000, Cell Signaling Technology, USA), rabbit anti-TrkB (1:1000, Cell Signaling Technology, USA), rabbit anti-PSD-95 (1:5000, Proteintech, USA), rabbit anti-SYN (1:1000, Cell Signaling Technology, USA), rabbit anti-SAP97 (1:1000, Cell Signaling Technology, USA), and mouse anti-tubulin (1:1000, Cell Signaling Technology, USA) were incubated overnight at 4 LJ. Subsequently, membranes were incubated with the respective secondary antibodies for 1 hour at room temperature and analyzed using a Bio-Rad ChemiDoc MP Chemiluminescence Gel Imaging System. GAPDH and tubulin were utilized as the internal standard.

## Statistical analyses

The statistical analysis was conducted using SPSS 26.0 software (SPSS, Chicago, IL, USA). The Kolmogorov-Smirnov test was employed to assess the normality of the distribution. Variations in the scale variable among groups were examined using two-way analysis of variance (ANOVA), succeeded by Tukey’s test for multiple comparisons, or by the Kruskal-Wallis test, accompanied by Dunn’s post hoc test. We also evaluated the treatment by age group interaction followed by analysis of simple effects. Body weights were evaluated using repeated-measures ANOVA, incorporating group and time as factors, followed by Tukey’s post-hoc testing. Quantitative data were presented as mean ± standard deviation (SD). Figures were created using GraphPad Prism 10.0 software (GraphPad Software Inc., San Diego, CA, USA). Image J software was employed for image processing and analysis in immunohistochemistry, hematoxylin and eosin staining, Golgi-Cox staining, and Western blotting. The Case Viewer 2.4 program was utilized to examine and take images. P-values below 0.05, 0.01, and 0.001 were regarded as thresholds for statistically significant differences.

## 3 Results

### 3.1 The 12 months old FTO-KD mice are more susceptible to CUMS-induced depression-like behaviors and cognitive dysfunction

Figure 2A illustrates that the sham group experienced a consistent increase in body weight in 3-and 12-month-old mice, whereas the CUMS group experienced a minor decrease in body weight. It is important to note that the body weight difference between the sham group and CUMS group was significant (P<0.001) in the sixth week. Additionally, the FTO-KD treatment substantially reduced the body weight of mice in the CUMS group in the 3-and 12-month-old groups (P<0.001 and P<0.05).

**Figure 2.**
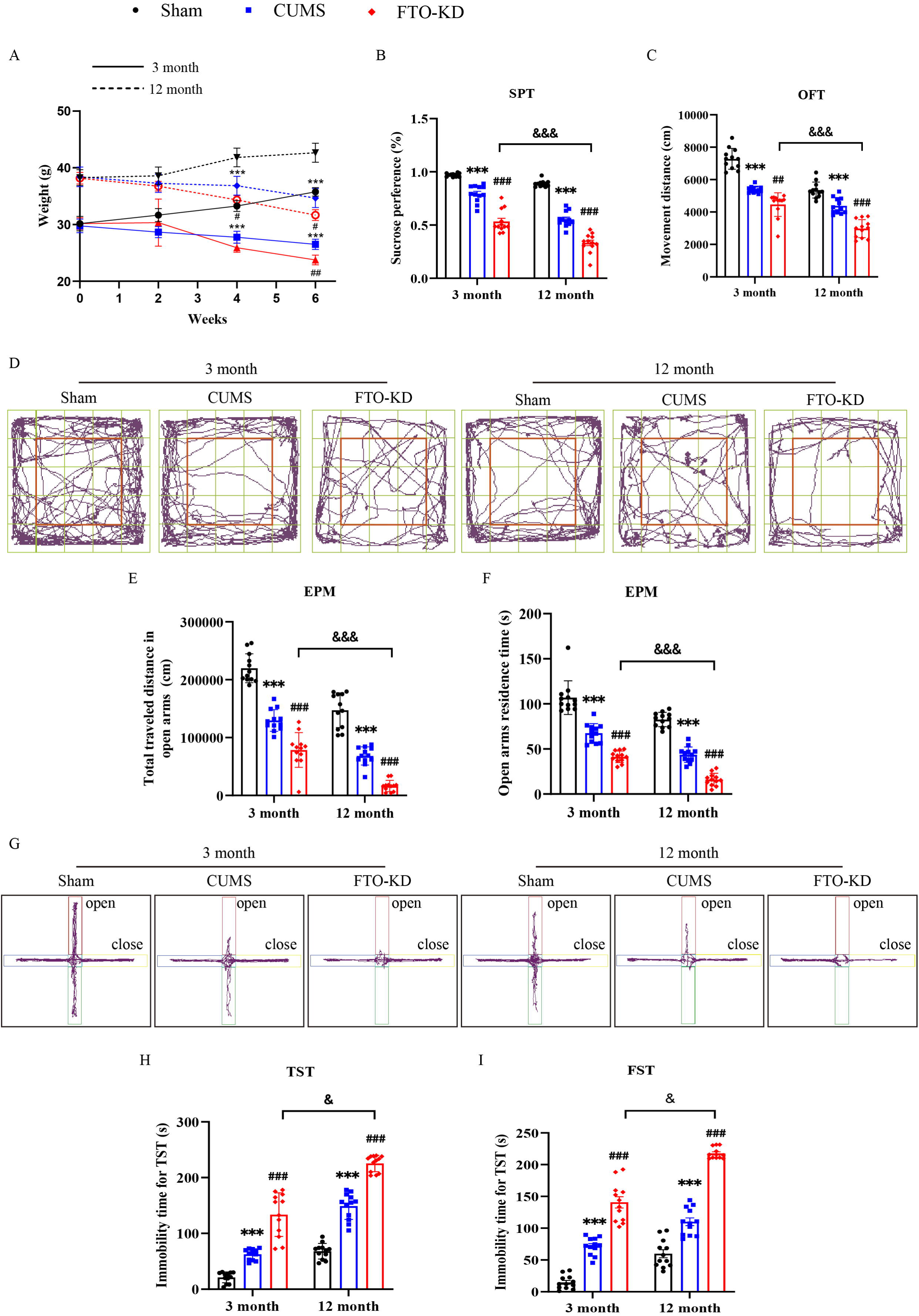
FTO deficiency aggravates CUMS-induced depression-like and anxiety-like behavior in an age-dependent manner. **A**. Weight changes in 3-month-old and 12-month-old mice. **B**. Sucrose preference percentage in SPT. **C**. Movement distance (cm) in OFT. **D**. Representative picture of the movement trajectory in OFT. **E**. The traveled distance in open arms. **F**. The time of residence in the open arms. **G**. Representative picture of the movement trajectory in EPM. **H**. Immobility time (s) in TST. **I**. Immobility time (s) in FST. N=12 per group. CUMS, Chronic Unpredictable Mild Stress; FTO-KD, FTO knockdown; SPT, Sucrose preference test; OFT, open field test; EPM, the elevated plus maze; TST, tail suspension test; FST, forced swimming test. Date was expressed as mean ± SD and were analyzed by two-way ANOVA followed by Tukey’s multiple comparisons post hoc test. \**P*<0.05, Sham vs. CUMS; ^#^*P*<0.05, CUMS vs FTO-KD; ^&^*P*<0.05, FTO-KD (3-month-old) vs. FTO-KD (12-month-old).

The behavioral experiments were conducted to assess the impact of FTO on depression-like and anxiety-like behaviors in CUMS mice of varying ages. Our findings indicated that the CUMS group exhibited a significantly lower sucrose preference (%) in SPT (P<0.001), movement distance in OFT (P<0.001), total traveled distance in open arms (P<0.001), and open arms residence time (P<0.001) in EPM compared to the age-corresponding sham group in the 3-and 12-month categories (Figure 2B-G).

Figure 2D and figure 2G are representative images of the motion trajectories of OFT and EPM, respectively. The CUMS group exhibited a significant increase in immobility time (TST, P<0.001; FST, P<0.001) in comparison to the corresponding placebo group at the 3-and 12-month groups (Figure 2H-I). These findings suggested that CUMS can induce weight loss, anhedonia, decreased activity, desperate behavior, and anxiety-like behavior in mice. The FTO-KD intervention exacerbated the aforementioned behaviors, thereby aggravating the depression-like and anxiety-like behaviors induced by CUMS in the 3-and 12-month-old groups. The behaviors induced by FTO-KD were significantly more severe in aged mice than in juvenile mice.

The MWM was implemented to evaluate the impact of FTO on neurocognitive function in CUMS mice. In the 3-and 12-month-old groups, the number of entries in the platform, time spent in the target quadrant, and number of entries in the target quadrant were substantially reduced in the CUMS group compared to the corresponding sham group, as illustrated in Figure 3. The aforementioned indicators were further diminished by FTO-KD, and the reduction effect was more pronounced in the 12-month-old group than in the 3-month-old group. The findings indicated that FTO-KD exacerbated the spatial memory impairment induced by CUMS, and older rodents appeared to be more susceptible. The locomotor activity of rodents in water was not influenced by FTO-KD.

**Figure 3.**
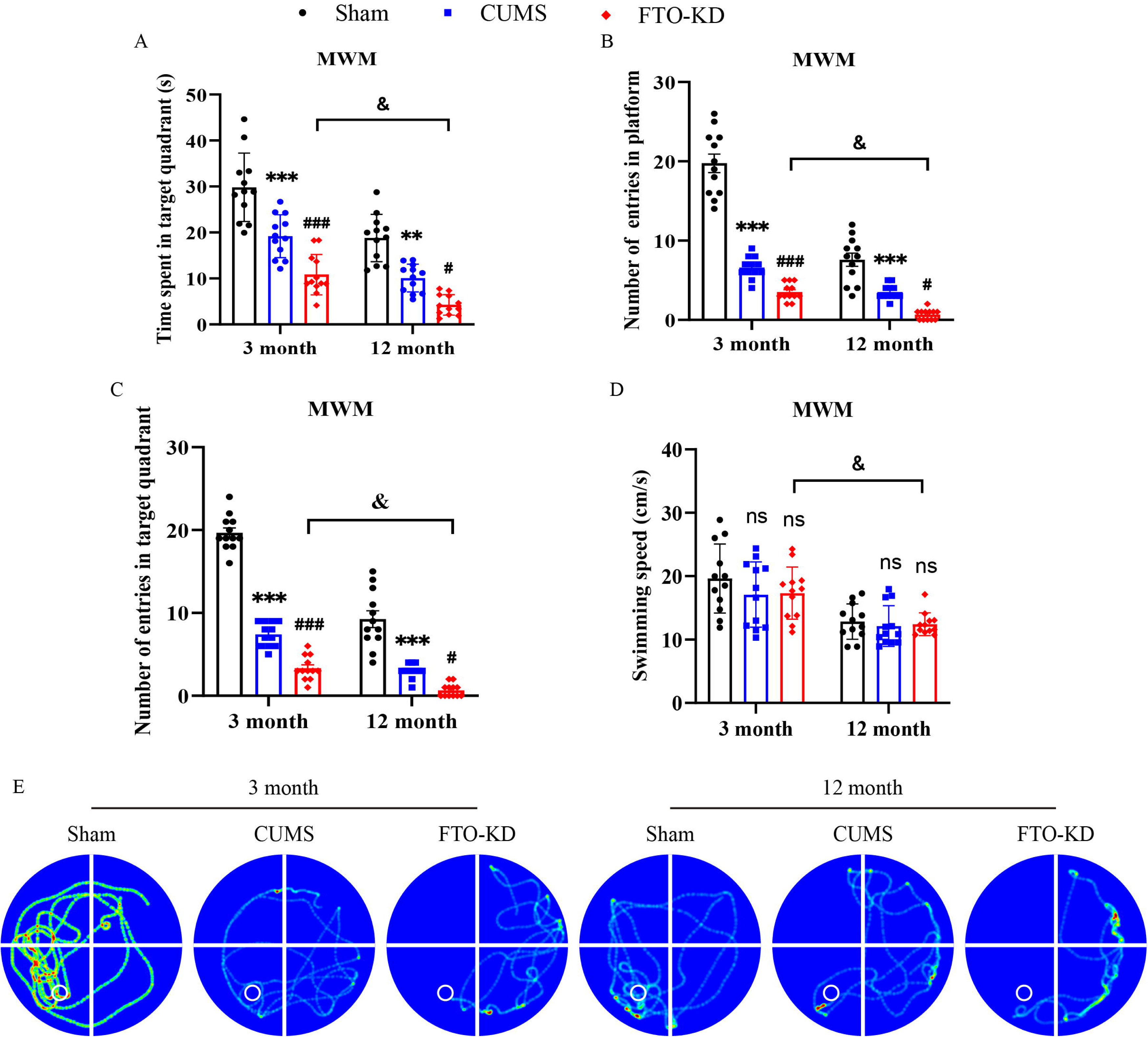
FTO deficiency aggravates CUMS-induced cognitive impairment in Morris Water Maze. **A.** Time spent in target quadrant (s); **B**. number of entries in platform. **C**. number of entries in target quadrant. **D**. Swimming speed (cm/s). E. Representative picture of the movement trajectory in MWM. N=12 per group. CUMS, Chronic Unpredictable Mild Stress; FTO-KD, FTO knockdown; MWM, Morris Water Maze. Date was expressed as mean ± SD and were analyzed by two-way ANOVA followed by Tukey’s multiple comparisons post hoc test. \**P*<0.05, Sham vs. CUMS; ^#^*P*<0.05, CUMS vs FTO-KD; ^&^*P*<0.05, FTO-KD (3-month-old) vs. FTO-KD (12-month-old).

Additionally, we identified a significant interaction pattern between the treatment and age categories on the aforementioned behaviors (**Supplementary Materials 1**). The behavioral differences among age groups were indicated by simple effect analysis (**Supplementary Materials 3**).

### 3.2 The expression of FTO protein in hippocampus of 12-month-old mice was significantly reduced after FTO-KD

Western blotting and IHC were employed to investigate the transfection of hippocampal FTO-KD AAV9 and its impact on hippocampal FTO expression. The expression of FTO in the hippocampus of mice in the CUMS group was significantly reduced in the 3-month-old (P<0.001) and 12-month-old groups (P<0.01) compared to the corresponding sham group, as confirmed by Western blotting results. The protein expression of FTO in the hippocampus of rodents in the CUMS group was further reduced by the intervention of FTO-KD, and the difference was statistically significant (P<0.05 and P<0.01, Figure 4A, B).

**Figure 4.**
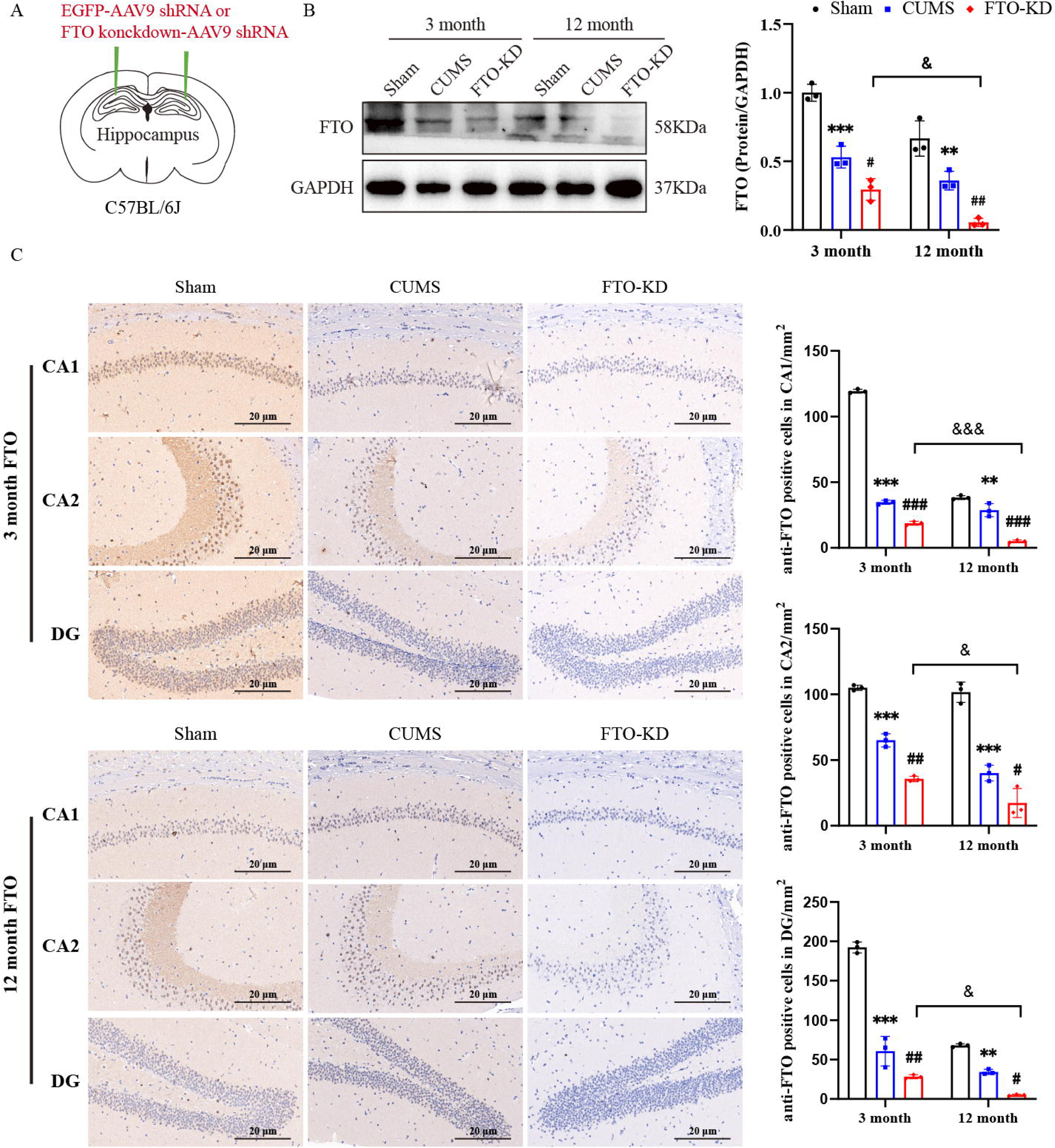
Brain stereotaxic surgery and expression of FTO in the hippocampus. **A.** FTO knockdown AAV9 shRNA virus injection. **B**. Representative Western blotting bands of FTO in the hippocampus. Quantification of OFT in hippocampus. **C.** Immunohistochemical staining of anti-FTO in the hippocampus. The anti-FTO-positive neurons were shown as dark brown dots. Scale bar = 20 µm. N=3 per group. CUMS, Chronic Unpredictable Mild Stress; FTO-KD, FTO knockdown. Date was expressed as mean ± SD and were analyzed by two-way ANOVA followed by Tukey’s multiple comparisons post hoc test. \**P*<0.05, Sham vs. CUMS; ^#^*P*<0.05, CUMS vs FTO-KD; ^&^*P*<0.05, FTO-KD (3-month-old) vs. FTO-KD (12-month-old).

In particular, the hippocampal FTO of 12-month-old rodents was more susceptible to FTO-KD. The IHC results, as illustrated in Figure 4C, indicated that the number of anti-FTO positive neurons in CA1, CA2, and DG of the CUMS group was substantially lower than that of the sham group at 3 and 12 months of age. The number of anti-FTO-positive neurons in the CA1, CA2, and DG regions of CUMS mice in all age groups was significantly diminished after FTO-KD intervention. The reduction was more pronounced in the 12-month-old group. These findings suggested that CUMS induces a decrease in FTO expression in the hippocampus of mice, FTO-KD is effectively integrated in the hippocampus, and older mice are more susceptible to FTO-KD AAV9.

Western blotting and IHC revealed an interaction pattern between the treatment and age groups on FTO protein expression (Supplementary Materials Table1). The analysis of simple effects revealed significant differences in FTO among both age groups (Supplementary Materials Table 3).

### 3.3 FTO-KD reduces NGF and reelin expression in hippocampus of 12-month-old CUMS mice

Early studies [40, 41] demonstrated a close relationship between hippocampal synaptic plasticity and NGF and reelin. We used IHC to detect the expression of NGF and reelin proteins in the hippocampus. IHC results showed that the number of anti-NGF-positive neurons in the DG region (P<0.001 and P<0.05) and the number of anti-reelin-positive neurons in CA2 mice (P<0.05) in the CUMS group was significantly lower than that in the corresponding sham group in the 3-and 12-month-old groups, respectively (Figure 5 and Figure 6). The number of anti-NGF positive cells in the DG region was suppressed even more with FTO-KD intervention (P<0.001 and P<0.05), as did the number of anti-reelin positive cells in the CA2 region (P<0.01 and P<0.05). In particular, mice in the 12-month-old group were more susceptible to FTO-KD interference.

**Figure 5.**
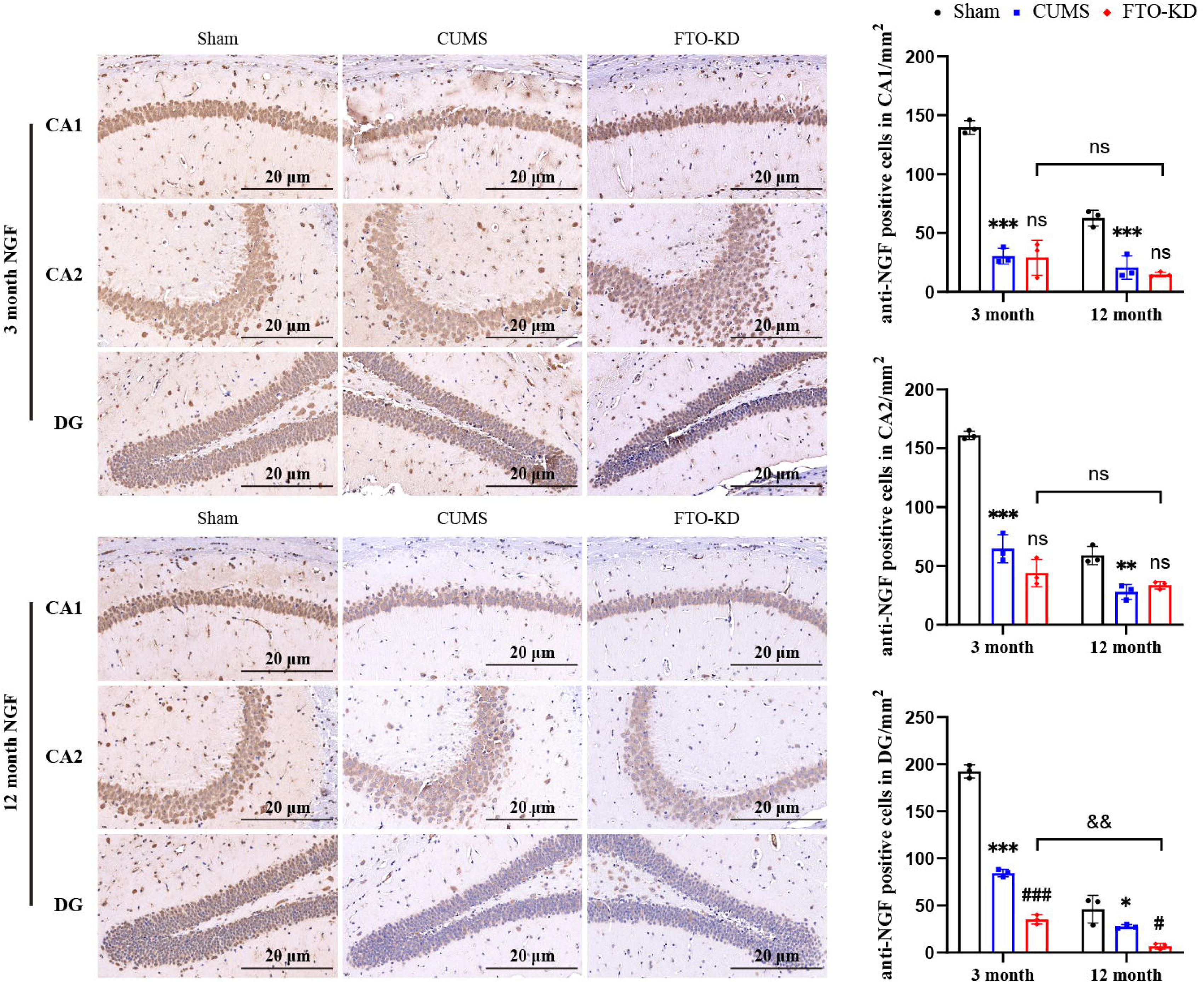
NGF expression in the hippocampus. Representative picture of NGF immunohistochemical staining in hippocampus. The anti-NGF-positive neurons were shown as dark brown dots. NGF-positive cells in CA1, CA2 and DG regions of the hippocampus were quantified. Scale bar = 20 µm. N=3 per group. CUMS, Chronic Unpredictable Mild Stress; FTO-KD, FTO knockdown. Date was expressed as mean ± SD and were analyzed by two-way ANOVA followed by Tukey’s multiple comparisons post hoc test. \**P*<0.05, Sham vs. CUMS; ^#^*P*<0.05, CUMS vs FTO-KD; ^&^*P*<0.05, FTO-KD (3-month-old) vs. FTO-KD (12-month-old).

**Figure 6.**
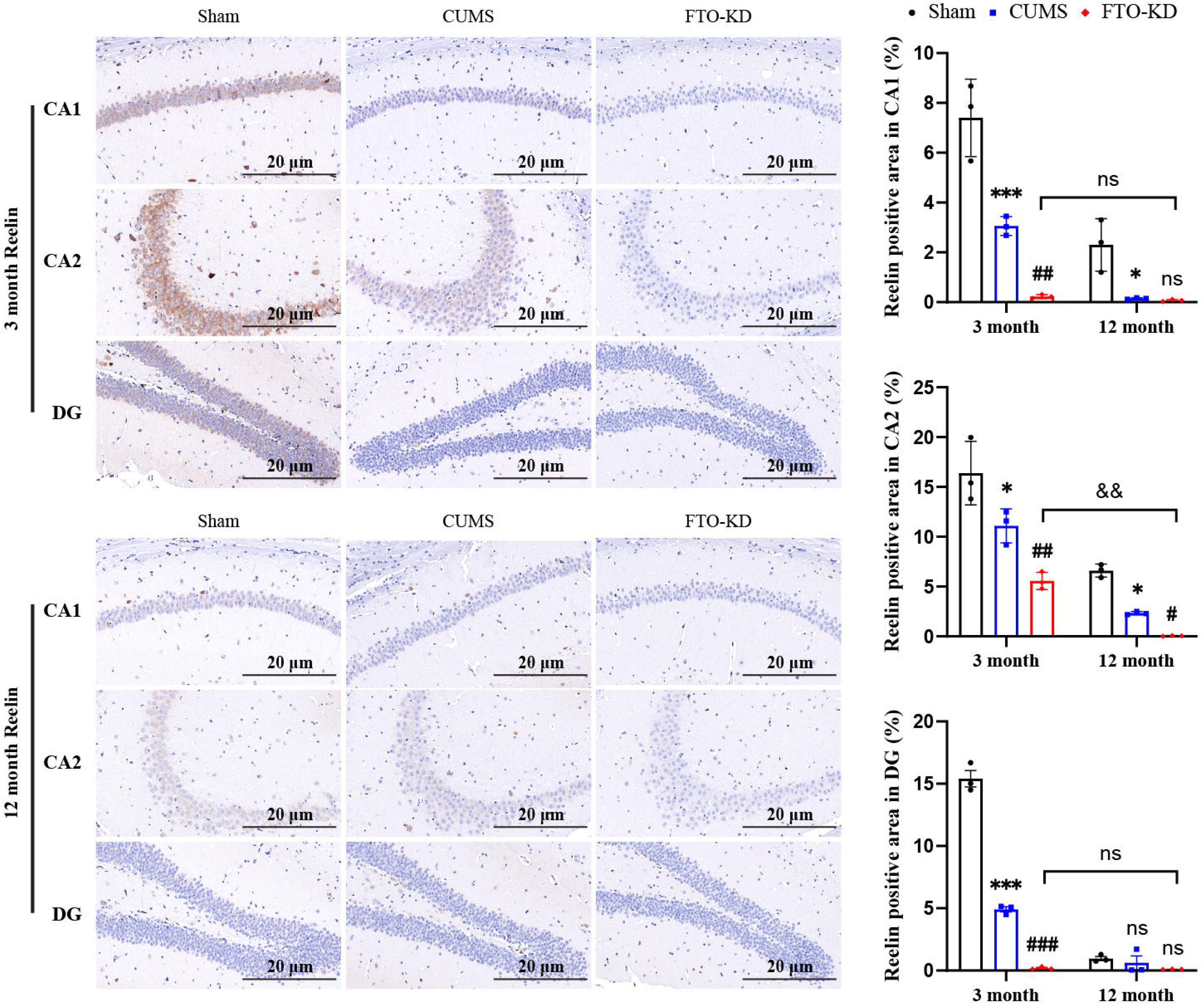
Reelin expression in the hippocampus. Representative picture of Reelin immunohistochemical staining in hippocampus. The anti-Reelin-positive neurons were shown as brown area. Reelin-positive cells in CA1, CA2 and DG regions of the hippocampus were quantified. Scale bar = 20 µm. N=3 per group. CUMS, Chronic Unpredictable Mild Stress; FTO-KD, FTO knockdown. Date was expressed as mean ± SD and were analyzed by two-way ANOVA followed by Tukey’s multiple comparisons post hoc test. \**P*<0.05, Sham vs. CUMS; ^#^*P*<0.05, CUMS vs FTO-KD; ^&^*P*<0.05, FTO-KD (3-month-old) vs. FTO-KD (12-month-old).

The interactions between treatment and age groups were significant in the CA1, CA2, and DG areas for NGF and reelin. A simple effects analysis showed that months of age changed the levels of NFG and reelin in different parts of the hippocampus, and these changes were statistically significant. Refer to Supplementary Materials Table 2 and Table 3 for further details.

Overall, CUMS may cause depressive-like behaviors and cognitive impairment in mice, possibly by lowering the expression of NGF and reelin proteins in the hippocampus. Furthermore, we highlight the connection between FTO loss and age in relation to cognitive impairment. FTO-KD exacerbated depression-like behaviors and cognitive deficits in an age-dependent manner.

### 3.4 FTO-KD aggravates hippocampal neuronal damage in 12-month-old CUMS mice

We used H&E staining to investigate the effect of FTO on hippocampal neurons. Figure 7 shows that in the hippocampal CA1 (P<0.05), CA2 (P<0.05), and DG (P<0.05 and P<0.01) areas, the CUMS group had more pyknotic nuclei than the sham group at both 3 and 12 months. FTO-KD intervention increased the number of pyknotic nuclei in hippocampal CA1 (*P*<0.01) and DG (*P*<0.01) areas of 12-month-old CUMS mice.

**Figure 7.**
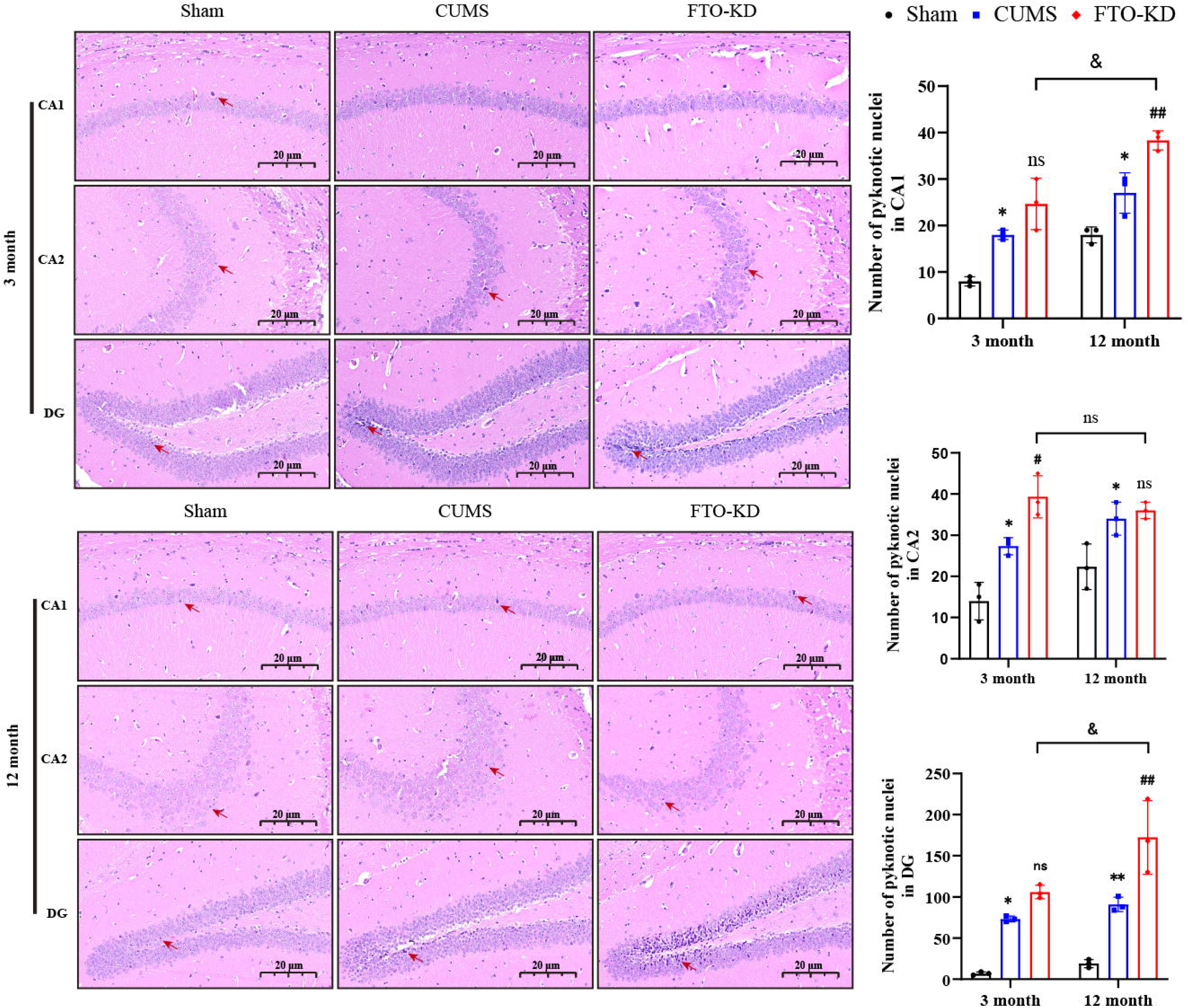
FTO deficiency aggravates CUMS-induced hippocampal neuronal damage. Representative image of H&E staining of the hippocampus. Quantification of neuronal pyknotic nuclei in CA1, CA2, and DG regions. Scale bar = 20 µm. N=3 per group. CUMS, Chronic Unpredictable Mild Stress; FTO-KD, FTO knockdown. Date was expressed as mean ± SD and were analyzed by two-way ANOVA followed by Tukey’s multiple comparisons post hoc test. \**P*<0.05, Sham vs. CUMS; ^#^*P*<0.05, CUMS vs FTO-KD; ^&^*P*<0.05, FTO-KD (3-month-old) vs. FTO-KD (12-month-old).

### 3.5 FTO-KD reduces hippocampal dendritic spine density in 12-month-old CUMS mice

Dendritic spines play an important role in neuronal plasticity [42]. We applied Golgi-cox staining to explore the impact of FTO on hippocampal synaptic plasticity. Using Golgi-cox staining, we found that the density of hippocampal dendritic spines was significantly reduced in the 3-and 12-month-old groups of CUMS mice compared to the corresponding sham group (*P*<0.05). Interestingly, the density of dendritic spines in 12-month-old rats was significantly lower than that in 3-month-old rats after FTO-KD intervention (Figure 8). These results showed that FTO-KD aggravated synaptic plasticity in the mouse hippocampus in an age-dependent manner.

**Figure 8.**
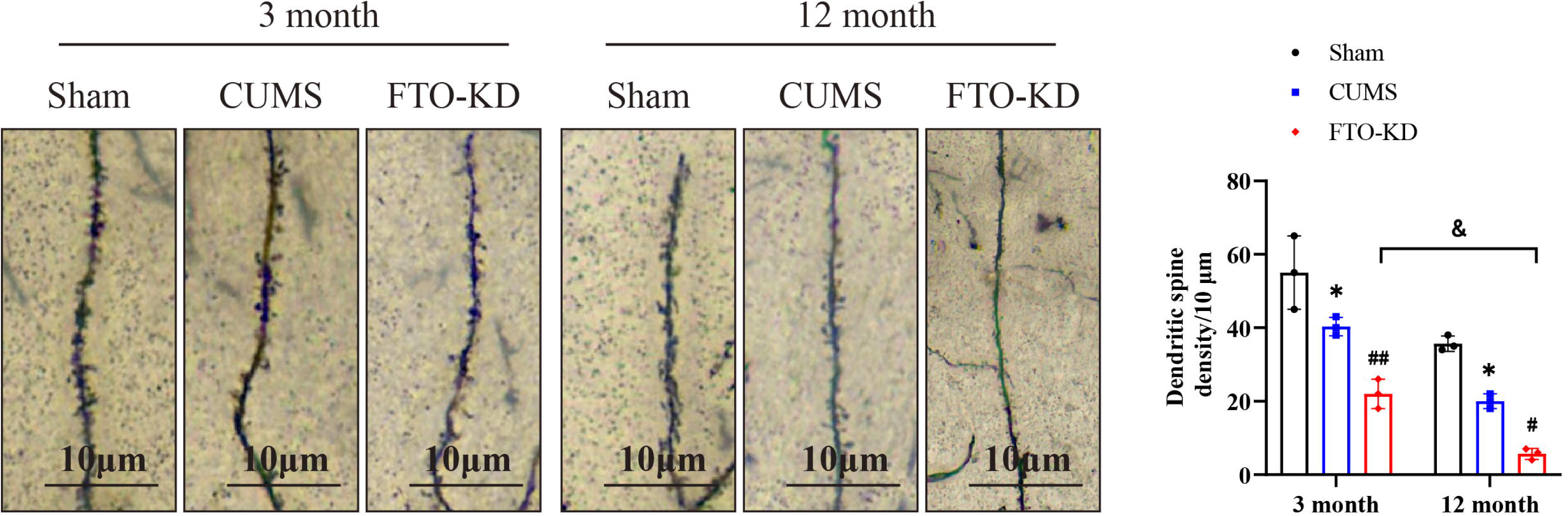
FTO deficiency aggravated CUMS-induced hippocampal dendritic spine abnormalities. Representative image of hippocampal dendritic spines. Quantification of hippocampal dendritic spine density. Scale bar = 10 µm. N=3 per group. CUMS, Chronic Unpredictable Mild Stress; FTO-KD, FTO knockdown. Date was expressed as mean ± SD and were analyzed by two-way ANOVA followed by Tukey’s multiple comparisons post hoc test. \**P*<0.05, Sham vs. CUMS; ^#^*P*<0.05, CUMS vs FTO-KD; ^&^*P*<0.05, FTO-KD (3-month-old) vs. FTO-KD (12-month-old).

### 3.6 FTO-KD attenuated the hippocampal expression of synaptic plasticity proteins in 12-month-old CUMS mice

Neurotrophins and synaptic plasticity proteins are directly involved in neuronal plasticity and play an important role in learning and memory [43]. In addition, evidence [44] suggests that the BDNF-TrkB signaling pathway regulates the development of neurons and the function of mature synapses. We used Western blotting and found that the levels of BNDF (P<0.001), TrkB (P<0.001 and P<0.05), PSD-95 (P<0.05), SYN (P<0.01), and SAP97 (P<0.01 and P<0.05) proteins were significantly lower in the CUMS group than in the sham group for both the 3-and 12-month-old groups.

Also, after FTO-KD intervention, the levels of BNDF, TrkB, PSD-95, SYN, and SAP97 proteins in the hippocampus of 12-month-old CUMS mice were lower than those in 3-month-old CUMS mice (Figure 9). The results showed that FTO-KD aggravated synaptic plasticity damage in the hippocampus by lowering the levels of BNDF, TrkB, PSD-95, SYN, and SAP proteins. In addition, 12-month-old CUMS mice seem to be more susceptible to FTO-KD. Supplementary Materials Table2 provides detailed data.

**Figure 9.**
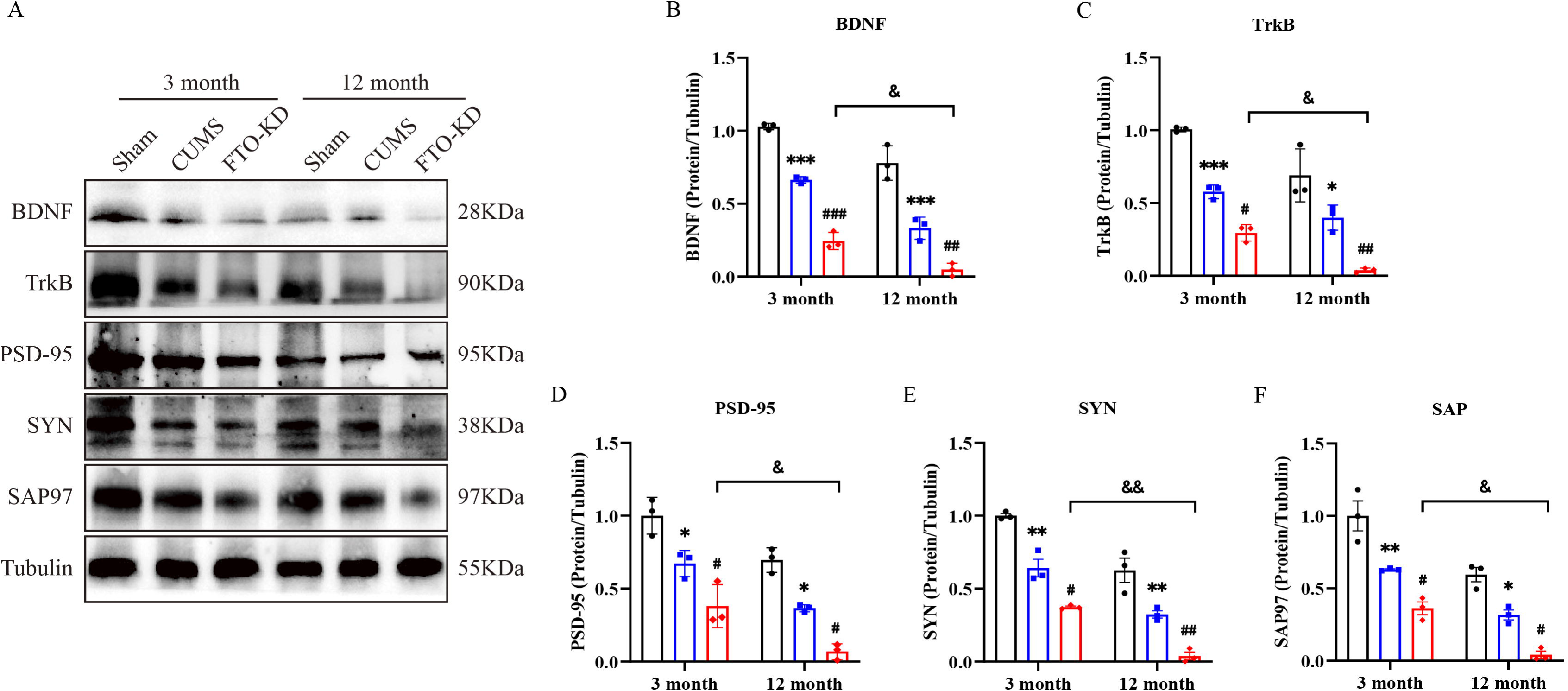
FTO deficiency aggravates synaptic plasticity damage in hippocampus. **A.** Representative image of Western blotting bands. **B-F**. Quantification of hippocampal BDNF, TrkB, PSD-95, SYN, and SAP97 expression. N=3 per group. CUMS, Chronic Unpredictable Mild Stress; FTO-KD, FTO knockdown; BDNF, brain-derived neurotrophic factor; TrkB, tyrosine Kinase receptor B; PSD-95, postsynaptic density protein 95; SYN, synaptophysin; SAP97, synapse associated protein of 97. Date was expressed as mean ± SD and were analyzed by two-way ANOVA followed by Tukey’s multiple comparisons post hoc test. \**P*<0.05, Sham vs. CUMS; ^#^*P*<0.05, CUMS vs FTO-KD; ^&^*P*<0.05, FTO-KD (3-month-old) vs. FTO-KD (12-month-old).

## 4. Discussion

### FTO-KD, age, and depression-and anxiety-like behaviors and cognitive deficits

This study effectively induced depression-like behavior and cognitive deficits in 3-and 12-month-old C57BL/6J mice following a 6-week CUMS paradigm. Notably, in comparison to the age-matched sham group, the 12-month-old mice exhibited a greater propensity for developing CUMS-induced depressive-like behaviors and cognitive impairments. Significantly, 12-month-old CUMS mice exhibited greater susceptibility to FTO than 3-month-old CUMS mice, and FTO deficiency exacerbated depressive-like behavior and cognitive impairment more in older CUMS mice than in their younger counterparts. Moreover, relative to the age-matched sham group, the expression of FTO protein in the hippocampus of CUMS group mice was markedly diminished, and FTO-KD AAV9 further attenuated the FTO expression levels in the hippocampus of CUMS mice, demonstrating that FTO-KD AAV9 effectively integrated into the hippocampus.

We found that the CUMS paradigm markedly decreased the expression levels of NGF and reelin in the hippocampus, leading to neuronal nuclear atrophy, disorganized neuronal architecture, diminished dendritic spine density, impaired BDNF-TrkB signaling, and reduced expression of synaptic plasticity-related proteins (PSD-95, SYN, and SAP97) in 3-and 12-month-old mice. FTO deficiency exacerbated hippocampal neuronal loss and pathological alterations in the hippocampus of CUMS animals, particularly in mice aged 12 months. The findings indicate that FTO deficiency exacerbates hippocampus neuronal dysfunction in CUMS animals in an age-dependent manner, resulting in depression-like behaviors and cognitive deficits. Consequently, FTO is a crucial facilitator of depressive-like behaviors in aged mice.

FTO variants correlate with food consumption and energy expenditure [39]. Our investigation revealed that FTO-KD markedly decreased the body weight of CUMS mice in both the 3-month and 12-month age groups. The reason may be that FTO inhibition activates STAT3 via ERK1/2, hence decreasing food consumption and body weight [40]. Results from SPT, OFT, TST, and FST demonstrated that CUMS exposure significantly diminished sucrose preference (%) and locomotor activity while augmenting immobility duration in 3-and 12-month-old mice compared to the age-matched sham group. Nonetheless, 12-month-old mice exhibited lower performance compared to 3-month-old mice in these tests. This suggests that the detrimental effects of CUMS in murine models are significantly more evident in older mice, thereby potentially heightening susceptibility to depression-like behaviors, consistent with previous studies [41, 42]. Concurrently, there is accumulating evidence that aging in rodents heightens their vulnerability to anhedonia [43].

Furthermore, we observed that FTO deficiency exacerbated CUMS-induced anhedonia, diminished exploratory activity, and increased desperate behavior in the SPT, OFT, TST, and FST behavioral assessments, particularly in aged mice. The findings indicate that FTO deletion heightens vulnerability to stress-induced depression-like behaviors in an age-dependent fashion, confirming that FTO downregulation markedly elevates the likelihood of developing depressive-like behaviors, particularly in older populations. Consequently, the cumulative effects of FTO deficiency and aging may serve as catalysts for the onset of depression-like behavior and cognitive impairment due to prolonged stress. It was previously shown that FTO knockout or knockdown can induce depression-like and anxiety-like behaviors in mice [44], as these mice exhibit increased anhedonia, reduced activity, and prolonged immobility time [45]. On the other hand, the overexpression of FTO or the introduction of exogenous FTO mitigated stress-induced depression-like behaviors [46].

Anxiety is among the most prevalent emotional symptoms in individuals with MDD. The World Health Organization reports that 4% of the global population today experiences anxiety [47]. Spychala et al. [48] discovered that FTO deficiency heightened anxiety and compromised long-term working memory in the hippocampus. Our work demonstrated that OFT and EPM revealed a substantial reduction in the crossing number, travel distance in open arms, and open arms residence time of mice following CUMS administration, indicating anxiety-like behaviors. The FTO-KD AAV9 intervention exacerbated anxiety-like behaviors triggered by CUMS exposure, aligning with prior findings. Nevertheless, several investigations have reported [49, 50] that FTO does not influence anxiety-like behaviors following acute or chronic stress in mice. These contradictory findings may be ascribed to differences in rodent species or the stress stimulation paradigm.

A recent study [51] indicated that naturally aging C57BL/6J mice exhibit cognitive decline from 6 months of age, with further deterioration observed at 22 months of age. In our study, 12-month-old CUMS mice exhibited significantly poorer performance than age-matched sham mice in the MWM. Furthermore, FTO-KD exacerbated the spatial learning and memory deficits caused by CUMS exposure, suggesting that FTO deletion heightened vulnerability to chronic stress-induced cognitive impairment in mice. Prior research [30, 52] has demonstrated that FTO deficiency can lead to reduced brain size in mice, alterations in brain region structures, and the inhibition of proliferation and differentiation of adult neural stem cells (aNSCs), resulting in compromised spatial learning and memory, aligning with our findings.

Polymorphisms in the FTO gene have been linked to a heightened incidence of depressive symptoms [53–55]. Nonetheless, there is conflicting information concerning the relationship between FTO and MDD. Preclinical findings [56] indicate that FTO deleted mice exhibit less resilience to stressors and a higher propensity for anxiety-like and depression-like behaviors. Conversely, another study [57] demonstrated that FTO whole-gene knockout mice lowered inflammatory factors via specific alterations in the gut microbiota, rendering the mice less vulnerable to stress stimuli and exhibiting diminished anxiety-like and depression-like behaviors. These differences may be due to a number of factors, including the heterogeneity of depression.

### FTO-KD, age and the hippocampus

Our study revealed that following CUMS exposure, FTO expression in the hippocampus of mice was diminished. Additionally, FTO-KD AAV9 treatment further decreased FTO expression in the hippocampus, exacerbating depressive-like behavior and cognitive deficits, with aged mice exhibiting increased vulnerability to chronic stress. The data unequivocally demonstrated that the loss of FTO following CUMS treatment adversely affects the hippocampus, with this damage exhibiting age-dependent characteristics. Furthermore, epigenetic modifications may have occurred with aging. In accordance with this, prior research [50] indicated that FTO expression was diminished in the hippocampus of individuals with significant depression and in animal models of depression. Exogenous FTO led to diminished m6A hypermethylation and ameliorated the depression-like phenotype [58].

There is growing evidence [59–61] that hippocampus neuronal plasticity, dendritic spine atrophy, and neurogenesis are significant factors in the pathophysiology of depression and cognitive impairment. Neurotrophins, including BDNF and NGF, play a role in synaptic plasticity [62], learning and memory [63], aging, neurogenesis [14], and neuronal maturation [64]. Furthermore, as a secreted glycoprotein, reelin is essential for neuronal development, dendritic growth and maturation, and synaptic plasticity [21]. Transcripts associated with BDNF are similarly implicated in synaptic plasticity inside the hippocampus, encompassing molecules pertinent to synaptic plasticity (PSD-95, SYN) and synaptic morphology (dendritic spine density and branching quantity). Studies [14, 65] indicates that mice subjected to prolonged unpredictable mild stress exhibit reduced expression levels of BDNF, NGF, reelin, PSD-95, and SYN in the hippocampus. Our findings indicate that NGF levels in the DG region, reelin in the CA2 region, dendritic spine density, and synaptic plasticity-related proteins such as PSD-95, SYN, and SAP97 were significantly diminished, while the activation of the BDNF-TrkB signaling pathway was aberrant in the hippocampus of mice subjected to CUMS at 3 and 12 months of age.

Furthermore, we observed pathological alterations including neuronal nucleus shrinkage and disordered neuronal cell shape in the hippocampus dentate gyrus region of the CUMS model. It is important to acknowledge that the interaction among the treatment and age groups significantly influenced the expression of both NGF and reelin proteins in the hippocampal subregions (CA1, CA2, and DG). Previous findings indicated that aberrant expression of these proteins and other pathological alterations in the hippocampus are prevalent neuropathological characteristics in patients with MDD and in animal models [66, 67].

Numerous studies indicate that FTO is integral to various biological processes, such as the regulation of neurogenesis [68], memory formation [69, 70], mood fluctuations [71], and synaptic plasticity [28]. FTO can modulate the expression of synaptic plasticity-related factors via its demethylase activity [72]. FTO may potentially regulate synaptic dysfunctions in hippocampus primary neurons by modulating synaptic activity and memory-associated mRNA [69, 73]. The present study revealed that FTO deficiency expedited the decline in hippocampal dendritic spine quantity, exacerbated neuronal disorganization and injury, and diminished the expression of proteins associated with synaptic plasticity, along with the aberrant transmission of the BDNF-TrkB signaling pathway in the CUMS model, particularly in aged mice. These findings reaffirm that FTO serves a significant neuroprotective function in depression. The absence of FTO results in depression-like behavior and cognitive deficits, as FTO deficiency induces m6A hypermethylation, causing the degradation of essential transcripts for synaptic plasticity (dendrites, BDNF, PSD-95, SYN, and SAP97) and neurogenesis (NGF, reelin), thereby downregulating their expression in the brain. Moreover, aging may serve as a trigger for neural deterioration. Wang et al. [73] similarly showed that the absence of FTO elevated m6A methylation levels of SYN mRNA and reduced SYN expression. Conversely, FTO overexpression reduced the m6A methylation level on SYN mRNA and elevated SYN expression. FTO, functioning as a demethylating agent of m6A in RNA, may be involved in synaptic plasticity by regulating the methylation modification of substrate mRNA.

## Limitation

This study has certain shortcomings that warrant acknowledgment. First, we did not conduct a reverse verification of the impact of FTO on depressive-like behavior and cognitive impairment in the CUMS model using exogenous FTO (FTO overexpression or FTO activator), particularly regarding neuroprotective benefits in aged mice. Second, this study utilized mice aged 3 and 12 months. Therefore, the effects of FTO in older (>12 months) mice has remained elusive. Third, this study only examined the impact of the demethylase FTO on depression, although other factors associated with m6A methylation, such as ALKBH5, WTAP, METTL3, and METTL14, are linked to mental depression. Fourth, our investigation focused solely on the influence of FTO in the hippocampus regarding depression-like behaviors, leaving other brain sub-regions (amygdala, prefrontal cortex, hypothalamus, striatum) unexplored. Fifth, sex-biased FTO targets have been documented to participate in brain development [74], although the present study does not include female mice.

## Conclusion

We discovered that 12-month-old mice exhibited greater susceptibility to CUMS and demonstrated inferior performance in depression-like and cognitive tasks relative to 3-month-old mice. Following FTO-KD treatment, the protein expression levels of FTO, NGF, and reelin in the hippocampus of 3-and 12-month-old CUMS mice were markedly diminished. Additionally, neuronal atrophy was pronounced, dendritic spine density was reduced, and the expression of proteins associated with synaptic plasticity was down-regulated, particularly in the 12-month-old cohort. This study reinforces the association between FTO and depression, elucidating the mechanism by which diminished FTO exacerbates age-related vulnerability to stress-induced depressive behaviors and cognitive deficits.

FTO deficiency and aging exacerbate vulnerability to depression-like behaviors and cognitive impairment, as well as hippocampus neuronal dysfunction. FTO is significant in the pathophysiology of MDD and the age-related vulnerability to develop MDD. The development of specific small molecule FTO activators may provide a novel therapeutic strategy for depression-like behaviors.

## Supporting information

ESF 1

## Abbreviations

FTO: Fat mass and obesity-related protein
CUMS: Chronic unpredictable mild stress
FTO-KD: AAV9 FTO knockdown adeno-associated virus 9 shRNA
NGF: Nerve growth factor
MDD: Major depressive disorder
BDNF: Brain-derived neurotrophic factor
PSD-95: Postsynaptic density protein 95
SYN: Synaptophysin
GABA: γ-aminobutyric acid
m6A: N6-methyladenosine
CNS: Central nervous system
EGFP: Enhanced green fluorescent protein
SPT: Sucrose preference test
OFT: Open field test
EPM: Elevated-plus maze
TST: Tail suspension test
FST: Forced swimming test
MWM: Morris Water Maze
IHC: Immunohistochemistry
H&E: Hematoxylin-eosin
ANOVA: Analysis of variance
SD: Standard deviation
aNSCs: Adult neural stem cells

## Supplementary information

Supplementary Materials Table1, 2, 3.

## Acknowledgements

Not applicable.

## Authors’ contributions

Original Draft Preparation, M.L; Methodology, Y.Z; Visualization, Y.Y, T.C, and Y.L; Review & Editing, M.M; Supervision, H.L and M.M; Project Administration, H.L and M.M; Funding Acquisition, H.L. All authors read and approved the final manuscript.

## Data availability

The dataset analyzed in this study is available from the corresponding author upon reasonable request.

## Declarations

### Ethics approval and consent to participate

The Ethics Committee of Chaohu Hospital affiliated to Anhui University of Medical Science approved all animal experiments (KYXM-202112-010).

### Consent for publication

Not applicable.

## Reference

1. Afridi MI, Hina M, Qureshi IS, Hussain M (2011) Cognitive disturbance comparison among drug-naive depressed cases and healthy controls. J Coll Physicians Surg Pak 21(6):351–355.

2. Story TJ, Potter GG, Attix DK, Welsh-Bohmer KA, Steffens DC (2008) Neurocognitive correlates of response to treatment in late-life depression. Am J Geriatr Psychiatry 16(9):752–759. 10.1097/JGP.0b013e31817e739a

3. Rock PL, Roiser JP, Riedel WJ, Blackwell AD (2014) Cognitive impairment in depression: a systematic review and meta-analysis. Psychol Med 44(10):2029–2040. 10.1017/S0033291713002535

4. Malhi GS, Mann JJ (2018) Depression. Lancet 392(10161):2299–2312. 10.1016/S0140-6736(18)31948-2

5. Bhalla RK, Butters MA, Mulsant BH, Begley AE, Zmuda MD, Schoderbek B, Pollock BG, Reynolds CF, 3rd, Becker JT (2006) Persistence of neuropsychologic deficits in the remitted state of late-life depression. Am J Geriatr Psychiatry 14(5):419–427. 10.1097/01.JGP.0000203130.45421.69

6. Mukku SSR, Dahale AB, Muniswamy NR, Muliyala KP, Sivakumar PT, Varghese M (2021) Geriatric Depression and Cognitive Impairment-An Update. Indian J Psychol Med 43(4):286–293. 10.1177/0253717620981556

7. Fenton EY, Fournier NM, Lussier AL, Romay-Tallon R, Caruncho HJ, Kalynchuk LE (2015) Imipramine protects against the deleterious effects of chronic corticosterone on depression-like behavior, hippocampal reelin expression, and neuronal maturation. Prog Neuropsychopharmacol Biol Psychiatry 60:52–59. 10.1016/j.pnpbp.2015.02.007

8. Jing C, Kong M, Ng KP, Xu L, Ma G, Ba M, Alzheimer’s Disease Neuroimaging I (2024) Hippocampal volume maximally modulates the relationship between subsyndromal symptomatic depression and cognitive impairment in non-demented older adults. J Affect Disord 367:640–646. 10.1016/j.jad.2024.09.018

9. Fan X, Wheatley EG, Villeda SA (2017) Mechanisms of Hippocampal Aging and the Potential for Rejuvenation. Annu Rev Neurosci 40:251–272. 10.1146/annurev-neuro-072116-031357

10. Numakawa T, Odaka H, Adachi N (2018) Actions of Brain-Derived Neurotrophin Factor in the Neurogenesis and Neuronal Function, and Its Involvement in the Pathophysiology of Brain Diseases. Int J Mol Sci 19(11). 10.3390/ijms19113650

11. Vilar M, Mira H (2016) Regulation of Neurogenesis by Neurotrophins during Adulthood: Expected and Unexpected Roles. Front Neurosci 10:26. 10.3389/fnins.2016.00026

12. Lu B, Pang PT, Woo NH (2005) The yin and yang of neurotrophin action. Nat Rev Neurosci 6(8):603–614. 10.1038/nrn1726

13. Ghosh A, Carnahan J, Greenberg ME (1994) Requirement for BDNF in activity-dependent survival of cortical neurons. Science 263(5153):1618–1623. 10.1126/science.7907431

14. Mondal AC, Fatima M (2019) Direct and indirect evidences of BDNF and NGF as key modulators in depression: role of antidepressants treatment. Int J Neurosci 129(3):283–296. 10.1080/00207454.2018.1527328

15. Martinez-Pinteno A, Mezquida G, Bioque M, Lopez-Ilundain JM, Andreu-Bernabeu A, Zorrilla I, Mane A, Rodriguez-Jimenez R, Corripio I, Sarro S et al (2022) The role of BDNF and NGF plasma levels in first-episode schizophrenia: A longitudinal study. Eur Neuropsychopharmacol 57:105–117. 10.1016/j.euroneuro.2022.02.003

16. Loch AA, Pinto MTC, Andrade JC, de Jesus LP, de Medeiros MW, Haddad NM, Bilt MTV, Talib LL, Gattaz WF (2023) Plasma levels of neurotrophin 4/5, NGF and pro-BDNF influence transition to mental disorders in a sample of individuals at ultra-high risk for psychosis. Psychiatry Res 327:115402. 10.1016/j.psychres.2023.115402

17. Filho CB, Jesse CR, Donato F, Giacomeli R, Del Fabbro L, da Silva Antunes M, de Gomes MG, Goes AT, Boeira SP, Prigol M et al (2015) Chronic unpredictable mild stress decreases BDNF and NGF levels and Na(+),K(+)-ATPase activity in the hippocampus and prefrontal cortex of mice: antidepressant effect of chrysin. Neuroscience 289:367–380. 10.1016/j.neuroscience.2014.12.048

18. Pesold C, Impagnatiello F, Pisu MG, Uzunov DP, Costa E, Guidotti A, Caruncho HJ (1998) Reelin is preferentially expressed in neurons synthesizing gamma-aminobutyric acid in cortex and hippocampus of adult rats. Proc Natl Acad Sci U S A 95(6):3221–3226. 10.1073/pnas.95.6.3221

19. D’Arcangelo G, Nakajima K, Miyata T, Ogawa M, Mikoshiba K, Curran T (1997) Reelin is a secreted glycoprotein recognized by the CR-50 monoclonal antibody. J Neurosci 17(1):23–31. 10.1523/JNEUROSCI.17-01-00023.1997

20. Fournier NM, Andersen DR, Botterill JJ, Sterner EY, Lussier AL, Caruncho HJ, Kalynchuk LE (2010) The effect of amygdala kindling on hippocampal neurogenesis coincides with decreased reelin and DISC1 expression in the adult dentate gyrus. Hippocampus 20(5):659–671. 10.1002/hipo.20653

21. Ibi D, Nakasai G, Sawahata M, Takaba R, Kinoshita M, Yamada K, Hiramatsu M (2023) Emotional behaviors as well as the hippocampal reelin expression in C57BL/6N male mice chronically treated with corticosterone. Pharmacol Biochem Behav 230:173617. 10.1016/j.pbb.2023.173617

22. Cui X, Meng J, Zhang S, Rao MK, Chen Y, Huang Y (2016) A hierarchical model for clustering m(6)A methylation peaks in MeRIP-seq data. BMC Genomics 17 Suppl 7(Suppl 7):520. 10.1186/s12864-016-2913-x

23. Meyer KD, Saletore Y, Zumbo P, Elemento O, Mason CE, Jaffrey SR (2012) Comprehensive analysis of mRNA methylation reveals enrichment in 3’ UTRs and near stop codons. Cell 149(7):1635–1646. 10.1016/j.cell.2012.05.003

24. Shafik AM, Zhang F, Guo Z, Dai Q, Pajdzik K, Li Y, Kang Y, Yao B, Wu H, He C et al (2021) N6-methyladenosine dynamics in neurodevelopment and aging, and its potential role in Alzheimer’s disease. Genome Biol 22(1):17. 10.1186/s13059-020-02249-z

25. Wen Y, Fu Z, Li J, Liu M, Wang X, Chen J, Chen Y, Wang H, Wen S, Zhang K et al (2024) Targeting m(6)A mRNA demethylase FTO alleviates manganese-induced cognitive memory deficits in mice. J Hazard Mater 476:134969. 10.1016/j.jhazmat.2024.134969

26. Yu J, Chen M, Huang H, Zhu J, Song H, Zhu J, Park J, Ji SJ (2018) Dynamic m6A modification regulates local translation of mRNA in axons. Nucleic Acids Res 46(3):1412–1423. 10.1093/nar/gkx1182

27. Hess ME, Hess S, Meyer KD, Verhagen LA, Koch L, Bronneke HS, Dietrich MO, Jordan SD, Saletore Y, Elemento O et al (2013) The fat mass and obesity associated gene (Fto) regulates activity of the dopaminergic midbrain circuitry. Nat Neurosci 16(8):1042–1048. 10.1038/nn.3449

28. Mitsuhashi H, Nagy C (2023) Potential Roles of m6A and FTO in Synaptic Connectivity and Major Depressive Disorder. Int J Mol Sci 24(7). 10.3390/ijms24076220

29. Chokkalla AK, Mehta SL, Vemuganti R (2020) Epitranscriptomic regulation by m(6)A RNA methylation in brain development and diseases. J Cereb Blood Flow Metab 40(12):2331–2349. 10.1177/0271678X20960033

30. Gao X, Shin YH, Li M, Wang F, Tong Q, Zhang P (2010) The fat mass and obesity associated gene FTO functions in the brain to regulate postnatal growth in mice. PLoS One 5(11):e14005. 10.1371/journal.pone.0014005

31. Boissel S, Reish O, Proulx K, Kawagoe-Takaki H, Sedgwick B, Yeo GS, Meyre D, Golzio C, Molinari F, Kadhom N et al (2009) Loss-of-function mutation in the dioxygenase-encoding FTO gene causes severe growth retardation and multiple malformations. American journal of human genetics 85(1):106–111. 10.1016/j.ajhg.2009.06.002

32. Song X, Wang W, Ding S, Liu X, Wang Y, Ma H (2021) Puerarin ameliorates depression-like behaviors of with chronic unpredictable mild stress mice by remodeling their gut microbiota. J Affect Disord 290:353–363. 10.1016/j.jad.2021.04.037

33. Ibi D, Fujiki Y, Koide N, Nakasai G, Takaba R, Hiramatsu M (2019) Paternal valproic acid exposure in mice triggers behavioral alterations in offspring. Neurotoxicol Teratol 76:106837. 10.1016/j.ntt.2019.106837

34. Yang C, Sui G, Li D, Wang L, Zhang S, Lei P, Chen Z, Wang F (2021) Exogenous IGF-1 alleviates depression-like behavior and hippocampal mitochondrial dysfunction in high-fat diet mice. Physiol Behav 229:113236. 10.1016/j.physbeh.2020.113236

35. Takaba R, Ibi D, Watanabe K, Hayakawa K, Nakasai G, Hiramatsu M (2022) Role of sirtuin1 in impairments of emotion-related behaviors in mice with chronic mild unpredictable stress during adolescence. Physiol Behav 257:113971. 10.1016/j.physbeh.2022.113971

36. Zhuo C, Tian H, Chen G, Ping J, Yang L, Li C, Zhang Q, Wang L, Ma X, Li R et al (2023) Low-dose lithium mono-and adjunctive therapies improve MK-801-induced cognitive impairment and schizophrenia-like behavior in mice -Evidence from altered prefrontal lobe Ca(2+) activity. J Affect Disord 337:128–142. 10.1016/j.jad.2023.05.069

37. Lin LY, Zhang J, Dai XM, Xiao NA, Wu XL, Wei Z, Fang WT, Zhu YG, Chen XC (2016) Early-life stress leads to impaired spatial learning and memory in middle-aged ApoE4-TR mice. Mol Neurodegener 11(1):51. 10.1186/s13024-016-0107-2

38. Pan B, Xu L, Weng J, Wang Y, Ji H, Han B, Zhu X, Liu Y (2022) Effects of icariin on alleviating schizophrenia-like symptoms by regulating the miR-144-3p/ATP1B2/mTOR signalling pathway. Neurosci Lett 791:136918. 10.1016/j.neulet.2022.136918

39. Cecil JE, Tavendale R, Watt P, Hetherington MM, Palmer CN (2008) An obesity-associated FTO gene variant and increased energy intake in children. N Engl J Med 359(24):2558–2566. 10.1056/NEJMoa0803839

40. Hu F, Yan HJ, Gao CX, Sun WW, Long YS (2023) Inhibition of Hypothalamic FTO Activates STAT3 Signal through ERK1/2 Associated with Reductions in Food Intake and Body Weight. Neuroendocrinology 113(1):80–91. 10.1159/000526752

41. Bachis A, Cruz MI, Nosheny RL, Mocchetti I (2008) Chronic unpredictable stress promotes neuronal apoptosis in the cerebral cortex. Neurosci Lett 442(2):104–108. 10.1016/j.neulet.2008.06.081

42. Ghaffari-Nasab A, Badalzadeh R, Mohaddes G, Alipour MR (2021) Young plasma administration mitigates depression-like behaviours in chronic mild stress-exposed aged rats by attenuating apoptosis in prefrontal cortex. Exp Physiol 106(7):1621–1630. 10.1113/EP089415

43. Malatynska E, Steinbusch HW, Redkozubova O, Bolkunov A, Kubatiev A, Yeritsyan NB, Vignisse J, Bachurin S, Strekalova T (2012) Anhedonic-like traits and lack of affective deficits in 18-month-old C57BL/6 mice: Implications for modeling elderly depression. Exp Gerontol 47(8):552–564. 10.1016/j.exger.2012.04.010

44. Jiang Y, Zhang T, Yang L, Du Z, Wang Q, Hou J, Liu Y, Song Q, Zhao J, Wu Y (2023) Downregulation of FTO in the hippocampus is associated with mental disorders induced by fear stress during pregnancy. Behav Brain Res 453:114598. 10.1016/j.bbr.2023.114598

45. Wang XL, Wei X, Yuan JJ, Mao YY, Wang ZY, Xing N, Gu HW, Lin CH, Wang WT, Zhang W et al (2022) Downregulation of Fat Mass and Obesity-Related Protein in the Anterior Cingulate Cortex Participates in Anxiety-and Depression-Like Behaviors Induced by Neuropathic Pain. Front Cell Neurosci 16:884296. 10.3389/fncel.2022.884296

46. Xu Z, Zhu X, Mu S, Fan R, Wang B, Gao W, Kang T (2024) FTO overexpression expedites wound healing and alleviates depression in burn rats through facilitating keratinocyte migration and angiogenesis via mediating TFPI-2 demethylation. Molecular and cellular biochemistry 479(2):325–335. 10.1007/s11010-023-04719-x

47. 47. WHO (2023) Anxiety-disorders. https://www.who.int/zh/#

48. Spychala A, Ruther U (2019) FTO affects hippocampal function by regulation of BDNF processing. PLoS One 14(2):e0211937. 10.1371/journal.pone.0211937

49. Engel M, Eggert C, Kaplick PM, Eder M, Roh S, Tietze L, Namendorf C, Arloth J, Weber P, Rex-Haffner M et al (2018) The Role of m(6)A/m-RNA Methylation in Stress Response Regulation. Neuron 99(2):389–403 e389. 10.1016/j.neuron.2018.07.009

50. Liu S, Xiu J, Zhu C, Meng K, Li C, Han R, Du T, Li L, Xu L, Liu R, et al (2021) Fat mass and obesity-associated protein regulates RNA methylation associated with depression-like behavior in mice. Nat Commun 12(1):6937.10.1038/s41467-021-27044-7

51. Yanai S, Endo S (2021) Functional Aging in Male C57BL/6J Mice Across the Life-Span: A Systematic Behavioral Analysis of Motor, Emotional, and Memory Function to Define an Aging Phenotype. Frontiers in aging neuroscience 13:697621. 10.3389/fnagi.2021.697621

52. Li L, Zang L, Zhang F, Chen J, Shen H, Shu L, Liang F, Feng C, Chen D, Tao H et al (2017) Fat mass and obesity-associated (FTO) protein regulates adult neurogenesis. Hum Mol Genet 26(13):2398–2411. 10.1093/hmg/ddx128

53. Mehrdad M, Eftekhari MH, Jafari F, Nikbakht HA, Gholamalizadeh M (2021) Does vitamin D affect the association between FTO rs9939609 polymorphism and depression? Expert Rev Endocrinol Metab 16(2):87–93. 10.1080/17446651.2021.1889367

54. Rivera M, Locke AE, Corre T, Czamara D, Wolf C, Ching-Lopez A, Milaneschi Y, Kloiber S, Cohen-Woods S, Rucker J et al (2017) Interaction between the FTO gene, body mass index and depression: meta-analysis of 13701 individuals. Br J Psychiatry 211(2):70–76. 10.1192/bjp.bp.116.183475

55. Harbron J, van der Merwe L, Zaahl MG, Kotze MJ, Senekal M (2014) Fat mass and obesity-associated (FTO) gene polymorphisms are associated with physical activity, food intake, eating behaviors, psychological health, and modeled change in body mass index in overweight/obese Caucasian adults. Nutrients 6(8):3130–3152. 10.3390/nu6083130

56. Wu PF, Han QQ, Chen FF, Shen TT, Li YH, Cao Y, Chen JG, Wang F (2021) Erasing m(6)A-dependent transcription signature of stress-sensitive genes triggers antidepressant actions. Neurobiology of stress 15:100390. 10.1016/j.ynstr.2021.100390

57. Sun L, Ma L, Zhang H, Cao Y, Wang C, Hou N, Huang N, von Deneen KM, Zhao C, Shi Y, et al (2019) Fto Deficiency Reduces Anxiety-and Depression-Like Behaviors in Mice via Alterations in Gut Microbiota. Theranostics 9(3):721–733. 10.7150/thno.31562

58. Chang R, Huang Z, Zhao S, Zou J, Li Y, Tan S (2022) Emerging Roles of FTO in Neuropsychiatric Disorders. Biomed Res Int 2022:2677312. 10.1155/2022/267731

59. Xu X, Yan Y, Yang Z, Zhang T (2024) Down-regulation of RIPK3 prevents depression-like behaviors by restoring the synaptic plasticity and suppressing neuronal loss. J Affect Disord 365:213–221. 10.1016/j.jad.2024.08.088

60. Fang S, Wu Z, Guo Y, Zhu W, Wan C, Yuan N, Chen J, Hao W, Mo X, Guo X et al (2023) Roles of microglia in adult hippocampal neurogenesis in depression and their therapeutics. Front Immunol 14:1193053. 10.3389/fimmu.2023.1193053

61. Maynard KR, Hobbs JW, Rajpurohit SK, Martinowich K (2018) Electroconvulsive seizures influence dendritic spine morphology and BDNF expression in a neuroendocrine model of depression. Brain Stimul 11(4):856–859. 10.1016/j.brs.2018.04.003

62. Castren E, Antila H (2017) Neuronal plasticity and neurotrophic factors in drug responses. Mol Psychiatry 22(8):1085–1095. 10.1038/mp.2017.61

63. Tu Y, Han D, Liu Y, Hong D, Chen R (2024) Nicorandil attenuates cognitive impairment after traumatic brain injury via inhibiting oxidative stress and inflammation: Involvement of BDNF and NGF. Brain Behav 14(1):e3356. 10.1002/brb3.335

64. Morcuende S, Munoz-Hernandez R, Benitez-Temino B, Pastor AM, de la Cruz RR (2013) Neuroprotective effects of NGF, BDNF, NT-3 and GDNF on axotomized extraocular motoneurons in neonatal rats. Neuroscience 250:31–48. 10.1016/j.neuroscience.2013.06.050

65. Li P, Huang W, Chen Y, Aslam MS, Cheng W, Huang Y, Chen W, Huang Y, Wu X, Yan Y et al (2023) Acupuncture Alleviates CUMS-Induced Depression-Like Behaviors by Restoring Prefrontal Cortex Neuroplasticity. Neural plasticity 2023:1474841. 10.1155/2023/147484

66. 66. Amidfar M, Reus GZ, de Moura AB, Quevedo J, Kim YK (2021) The Role of Neurotrophic Factors in Pathophysiology of Major Depressive Disorder. Adv Exp Med Biol 1305:257–272. 10.1007/978-981-33-6044-0_14

67. Liu F, Jia Y, Zhao L, Xiao LN, Cheng X, Xiao Y, Zhang Y, Zhang Y, Yu H, Deng QE et al (2024) Escin ameliorates CUMS-induced depressive-like behavior via BDNF/TrkB/CREB and TLR4/MyD88/NF-kappaB signaling pathways in rats. Eur J Pharmacol 984:177063. 10.1016/j.ejphar.2024.177063

68. Gao H, Cheng X, Chen J, Ji C, Guo H, Qu W, Dong X, Chen Y, Ma L, Shu Q et al (2020) Fto-modulated lipid niche regulates adult neurogenesis through modulating adenosine metabolism. Hum Mol Genet 29(16):2775–2787. 10.1093/hmg/ddaa17

69. Widagdo J, Zhao QY, Kempen MJ, Tan MC, Ratnu VS, Wei W, Leighton L, Spadaro PA, Edson J, Anggono V et al (2016) Experience-Dependent Accumulation of N6-Methyladenosine in the Prefrontal Cortex Is Associated with Memory Processes in Mice. J Neurosci 36(25):6771–6777. 10.1523/JNEUROSCI.4053-15.2016

70. Walters BJ, Mercaldo V, Gillon CJ, Yip M, Neve RL, Boyce FM, Frankland PW, Josselyn SA (2017) The Role of The RNA Demethylase FTO (Fat Mass and Obesity-Associated) and mRNA Methylation in Hippocampal Memory Formation. Neuropsychopharmacology 42(7):1502–1510. 10.1038/npp.2017.31

71. Luppino FS, de Wit LM, Bouvy PF, Stijnen T, Cuijpers P, Penninx BW, Zitman FG (2010) Overweight, obesity, and depression: a systematic review and meta-analysis of longitudinal studies. Arch Gen Psychiatry 67(3):220–229. 10.1001/archgenpsychiatry.2010.2

72. Lin L, Hales CM, Garber K, Jin P (2014) Fat mass and obesity-associated (FTO) protein interacts with CaMKII and modulates the activity of CREB signaling pathway. Hum Mol Genet 23(12):3299–3306. 10.1093/hmg/ddu043

73. Wang Y, Wu Z, He Y, Zeng X, Gu Z, Zhou X, Si W, Chen D (2024) Fat mass and obesity-associated protein regulates RNA methylation associated with spatial cognitive dysfunction after chronic cerebral hypoperfusion. Neuropeptides 105:102428. 10.1016/j.npep.2024.102428

74. Chokkalla AK, Jeong S, Mehta SL, Davis CK, Morris-Blanco KC, Bathula S, Qureshi SS, Vemuganti R (2023) Cerebroprotective Role of N(6)-Methyladenosine Demethylase FTO (Fat Mass and Obesity-Associated Protein) After Experimental Stroke. Stroke 54(1):245–254. 10.1161/STROKEAHA.122.040401

